# Integrated analysis of single cell transcriptomic data across conditions, technologies, and species

**DOI:** 10.1101/164889

**Authors:** Andrew Butler, Rahul Satija

## Abstract

Single cell RNA-seq (scRNA-seq) has emerged as a transformative tool to discover and define cellular phenotypes. While computational scRNA-seq methods are currently well suited for experiments representing a single condition, technology, or species, analyzing multiple datasets simultaneously raises new challenges. In particular, traditional analytical workflows struggle to align subpopulations that are present across datasets, limiting the possibility for integrated or comparative analysis. Here, we introduce a new computational strategy for scRNA-seq alignment, utilizing common sources of variation to identify shared subpopulations between datasets as part of our R toolkit Seurat. We demonstrate our approach by aligning scRNA-seq datasets of PBMCs under resting and stimulated conditions, hematopoietic progenitors sequenced across two profiling technologies, and pancreatic cell ‘atlases’ generated from human and mouse islets. In each case, we learn distinct or transitional cell states jointly across datasets, and can identify subpopulations that could not be detected by analyzing datasets independently. We anticipate that these methods will serve not only to correct for batch or technology-dependent effects, but also to facilitate general comparisons of scRNA-seq datasets, potentially deepening our understanding of how distinct cell states respond to perturbation, disease, and evolution.

**Availability:** Installation instructions, documentation, and tutorials are available at http://www.satijalab.org/seurat

## INTRODUCTION

With recent improvements in cost and throughput^1-3^, accompanied by the availability of fully commercialized workflows^4^, high-throughput single cell transcriptomics has become a routine and powerful tool for unbiased profiling of complex and heterogeneous systems. In concert with novel computational approaches, these datasets can be used for unsupervised discovery of cell types and states^56^, the reconstruction of developmental trajectories and fate decisions^78^, and to spatially model complex tissues^9,10^. Indeed, single cell RNA-seq (scRNA-seq) is poised to transform our understanding of developmental biology and gene regulation^11-14^, and enable systematic reconstruction of cellular taxonomies across the human body^6,15^, though significant computational challenges remain.

In particular, there are no existing methods that enable integrated or comparative analysis of different scRNA-seq datasets consisting of multiple transcriptomic subpopulations, either to compare heterogeneous tissues across different conditions, or to integrate measurements produced by different technologies. Many powerful methods address individual components of this crucial challenge. For example, zero-inflated differential expression tests have been tailored to scRNA-seq data to identify changes within a single cell type^16,17^, and clustering approaches^18-23^ can detect proportional shifts across conditions if cell types are conserved. However, comparative analysis for scRNA-seq poses a unique challenge, as changes in the proportional composition of cell types in a sample are blended with expression changes within a given cell type, and simultaneous analysis of multiple datasets will confound these two disparate effects. Therefore, new methods are needed that can learn jointly between multiple datasets, for example, identifying and aligning a shared set of cell types that are present in both experiments, focusing initially on shared similarities to facilitate comparative analysis downstream.

Progress towards this goal is essential for translating the oncoming wealth of single cell sequencing data into biological insight. An integrated computational framework for joint learning between datasets would allow for robust and insightful comparisons of heterogeneous tissues in health and disease, for example, resolving cell-type specific responses to infection by comparative scRNA-seq of peripheral blood mononuclear cells (PBMCs). Crucially, these methods would be capable of integrating data from diverse technologies, potentially combining deep plate-based sequencing of single cells with sparse droplet technologies. In principle, they could also be used to identify and align cell types from distinct species, enabling evolutionary comparisons, and the discovery of convergent and divergent transcriptional programs.

Here, we present a novel computational strategy for integrated analysis of single cell RNA-seq datasets. We demonstrate that multivariate methods designed for ‘manifold alignment’^24,25^ can be successfully applied to scRNA-seq data, in order to identify gene-gene correlation patterns that are conserved across datasets, and embed cells in a shared low-dimensional space. We demonstrate the generality of our approach on diverse datasets from the literature with three distinct analyses: we identify and compare 13 aligned PBMC subpopulations under resting and IFN-β stimulated conditions, we integrate deeply sequenced SMART-Seq2 and sparse MARS-Seq data from murine bone marrow, and we jointly learn shared cell types from droplet-based ‘atlases’ of human and mouse pancreatic tissue. These analyses pose distinct challenges for alignment, but in each case we can successfully integrate the datasets, and learn deeper biological insight than would be possible from individual analysis. Our approach can be successfully applied to datasets ranging from hundreds to tens of thousands of cells, is compatible with diverse profiling technologies, and is implemented as part of Seurat, an open-source R toolkit for single cell genomics.

## RESULTS

We aimed to develop a diverse integration strategy that could compare scRNA-seq datasets across different conditions, technologies, or species. To be successful in diverse settings, this computational strategy must fulfill the following requirements, illustrated with a toy example where heterogeneous scRNA-seq datasets are generated in the presence or absence of a drug (Figure 1A). First, subpopulations must be aligned even if each has a unique drug response. This key challenge lies outside of the scope of batch correction methods, which assume that confounding variables have uniform effects on all cells in a dataset. Second, the method must allow for changes in cellular density (shifts in subpopulation frequency) between conditions. Third, the method must be robust to changes in feature scale across conditions, allowing either global transcriptional shifts, or differences in normalization strategies between datasets produced with different technologies (i.e. UMI vs. FPKM). Lastly, the process should be unsupervised, with no requirement for pre-established sets of markers that can be used to match subpopulations.

**Figure 1.**
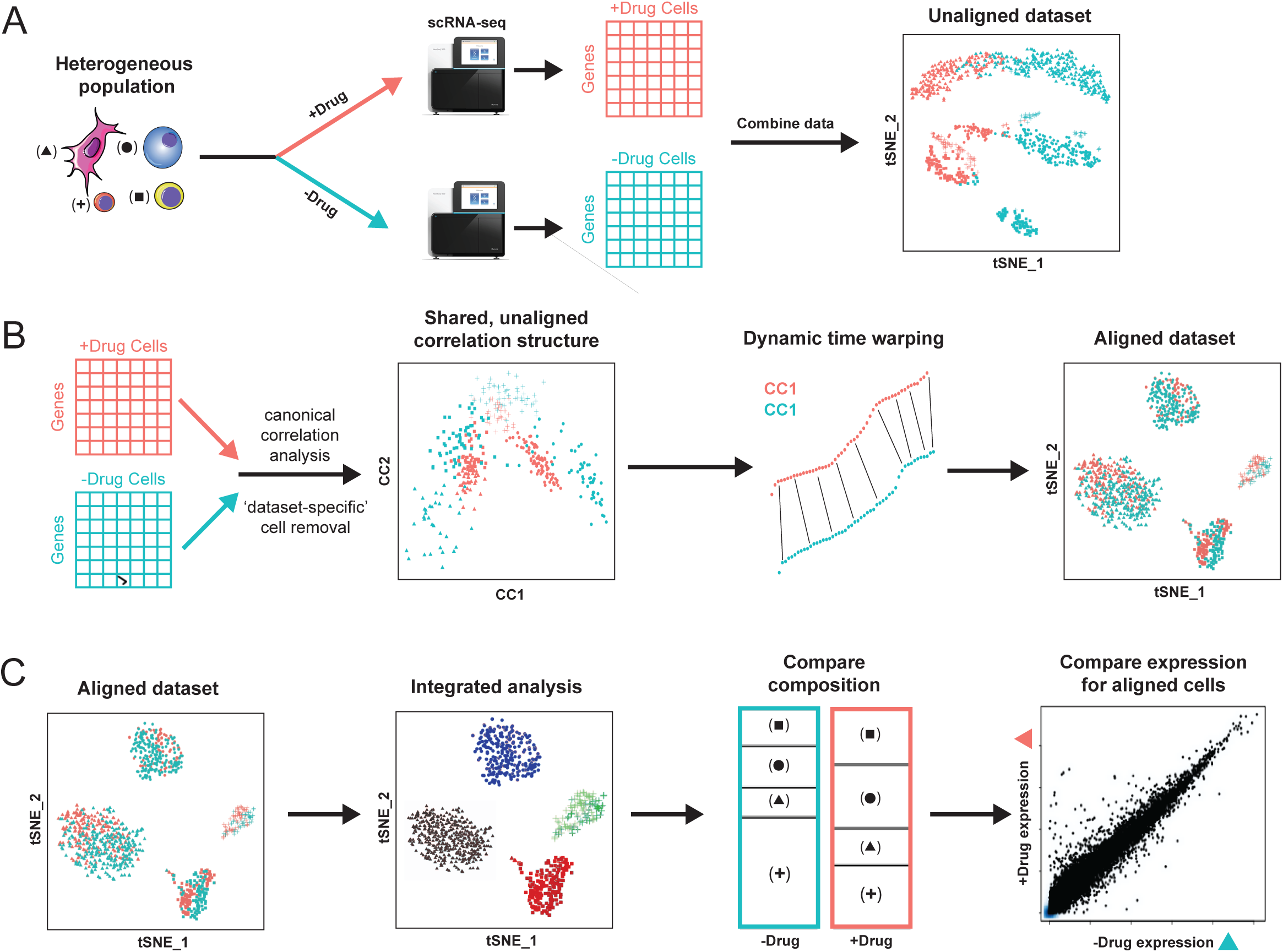
Overview of Seurat alignment of single cell RNA-seq datasets. **(A)** Toy example of heterogeneous populations profiled in a case/control study after drug treatment. Cells across four types are plotted with different s, while stimulation condition is encoded by color. In a standard workflow, cells often cluster both by cell type and stimulation condition, creating challenges for downstream comparative analysis. **(B)** The Seurat alignment procedure uses canonical correlation analysis to identify shared correlation structures across datasets, and aligns these dimensions using dynamic time warping. After alignment, cells are embedded in a shared low-dimensional space (visualized here in 2D with tSNE). **(C)** After alignment, a single integrated clustering can identify conserved cell types across conditions, allowing for comparative analysis to identify shifts in cell type proportion, as well as cell-type specific transcriptional responses to drug treatment.

The Seurat alignment workflow takes as input scRNA-seq data from two conditions, and briefly consists of the following steps (Figure 1B-C; Supplementary Methods). (i) It learns a shared gene correlation structure that is conserved between the two datasets, and can serve as a scaffold for the alignment (Figure 1B). (ii) It identifies and discards individual cells that cannot be well described by this shared structure, and are therefore ‘dataset-specific’ (unalignable). (iii) It aligns the two datasets into a conserved low-dimensional space, using non-linear ‘warping’ algorithms to normalize for differences in feature scale, in a manner that is robust to shifts in population density. (iv) It proceeds with an integrated downstream analysis, for example, identifying discrete subpopulations through clustering, or reconstructing continuous developmental processes (Figure 1C). (v) It performs comparative analysis on aligned subpopulations between the two datasets, to identify changes in population density or gene expression (Figure 1C). We describe each of these steps below, and then apply and validate this strategy on three dataset pairs in the literature.

### Identifying shared correlation structures across datasets

Machine-learning techniques for ‘data fusion’ aim to integrate information from multiple experiments into a consistent representation. For example, canonical correlation analysis (CCA) aims to find linear combinations of features across datasets that are maximally correlated, essentially, identifying shared correlation structures across datasets^26,27^. CCA has been used for multi-modal genomic analysis from bulk samples, for example identifying relationships between gene expression and DNA copy number measurements based on the same set of samples^28^. Here, in contrast to its traditional use in multi-modal analysis^29,30^, we apply CCA to identify relationships between single cells, from different datasets, based on the same set of genes. Effectively, we treat the datasets as multiple measurements of a gene-gene covariance structure, and search for patterns that are common to both datasets.

We employ a variant of CCA, diagonal CCA (Supplementary Methods), to account for cases where there are more cells than genes, and apply this using the two single-cell RNA-seq datasets as input. The procedure can consider any gene that is measured in both datasets, though we choose to focus only on genes that exhibit high single cell variation in either dataset independently (Supplementary Methods). CCA finds two sets of canonical ‘basis’ vectors, embedding cells from each dataset in a low-dimensional space, such that the variation along these vectors (gene-level projections) is highly correlated between datasets. We note that CCA is robust to affine transformations in the original data, and is unaffected by linear shifts in gene expression (for example, due to different normalization strategies). Additionally, CCA focuses on shared relationships between the data; if there are dataset-specific cell types that do not represent shared sources of variation between datasets, these cells should not be separated along the canonical basis vectors.

We leverage this to identify cells that cannot be aligned between the two datasets. Briefly, we quantify how well the low-dimensional space defined by CCA explains each cell’s expression profile, and compare this to PCA, which is performed on each dataset independently (Supplementary Methods). Cells where the percent variance explained is reduced by a user-defined cutoff in CCA compared to PCA are therefore defined by sources of variance that are not shared between the datasets. We use a cutoff of 50% for all examples here to identify these cells, and discard them from the alignment workflow.

### Aligning basis vectors from CCA

CCA returns vectors whose gene-level projections are correlated between datasets, but not necessarily aligned. While linear transformations may be required to correct for global shifts in feature scale or normalization strategy, non-linear shifts may also be needed to correct for shifts in population density. We therefore align the CCA basis vectors between the two datasets, resulting in a single, integrated low-dimensional space. Briefly, we represent each basis vector as a ‘metagene’, defined as a weighted expression average of the top genes whose expression exhibits robust correlations with the basis vector (Supplementary Methods). We first linearly transform the ‘metagenes’ to match their 95% reference range, correcting for global differences in feature scale. Next, we determine a mapping between the metagenes using ‘dynamic time warping’, which locally compresses or stretches the vectors during alignment to correct for changes in population density^31^. We apply this procedure to each pair of basis vectors individually, defining a single, aligned, low-dimensional space representing both datasets. This enables us to perform integrated downstream analyses, including unbiased clustering and the reconstruction of developmental trajectories, as demonstrated below.

### Comparative analysis of stimulated and resting PBMCs

We first demonstrate our alignment strategy on a dataset containing many distinct cell types in the presence and absence of perturbation. For example, a recent study examining the effects of interferon stimulation split 14,038 human PBMCs from 8 patients into two groups: one stimulated with interferon-beta (IFN-β) and a culture matched control^32^. Since all cells contain machinery to respond to IFN-β, stimulation results in a drastic but highly cell-type specific response. Consequently, a traditional joint analysis yields confusing results, as cells tend to cluster both by cell type but also by stimulation condition (Figure 2A). As an alternative to unbiased clustering, a supervised strategy to assign cells to classes based on known markers resulted in a final set of eight clusters^32^.

**Figure 2.**
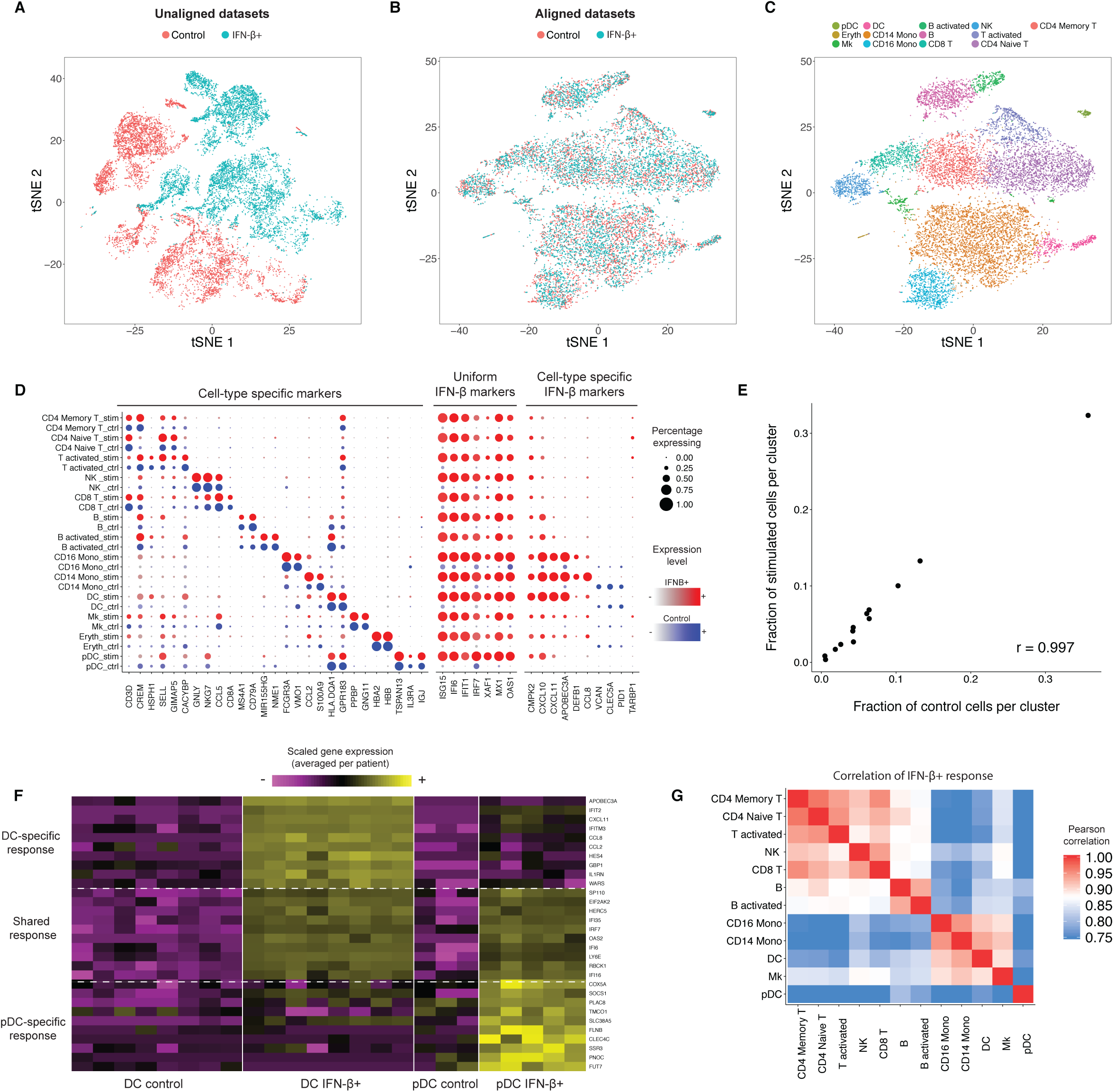
Integrated analysis of resting and stimulated PBMC. **(A-C)** tSNE plots of 14,038 human PBMCs split between control and IFN-β-stimulated conditions, prior to (A) and post (B) alignment. After alignment, cells across stimulation conditions group together based on shared cell type, allowing for a single joint clustering (C) to detect 13 immune populations. **(D)** Integrated analysis reveals markers of cell types (conserved across stimulation conditions), uniform markers of IFN-p+ response (independent of cell type), and components of the IFN-p+ response that vary across cell types. The size of each circle reflects the percentage of cells in a cluster where the gene is detected, and the color reflects the average expression level within each cluster. **(E)** The fraction of cells (median across 8 donors) falling in each cluster for stimulated and unstimulated cells. **(F)** Examples of heterogeneous responses to IFN-β between conventional and plasmacytoid dendritic cells (global analysis shown in Supplementary Figure 3B). Each column represents the average expression of single cells within a single patient. Only patient/cluster combinations with at least five cells are shown. **(G)** Correlation heatmap of cell-type specific responses to IFN-β (individual correlations for T and DC subsets shown in Supplementary Figure 3A-B). Cells from myeloid and lymphoid lineages show highly correlated responses, but plasmacytoid dendritic cells exhibit a unique IFN-β response.

In contrast, the Seurat alignment returned a set of canonical correlation vectors that separated PBMC subsets irrespective of stimulation condition. We performed joint graph-based clustering on these aligned vectors and visualized the results with t-Distributed Stochastic Neighbor Embedding (t-SNE) to verify that cells grouped entirely by cell type and were properly aligned across conditions (Figure 2B). Our analysis revealed 13 cell types, consistent with the published supervised analysis but with significantly greater resolution (Figure 2C; Supplementary Table 1). In particular, we were able to separate naïve from memory T cells, plasmacytoid dendritic cells (pDCs) from conventional dendritic cells, and identify an extremely rare (0.4%) population of contaminating erythroblasts. In addition, for T cells and B cells, we discovered activated subpopulations marked by a strong stress response expression signature that is likely an artifact of the culturing process in both conditions (Supplementary Figure 1A-B). We could verify the identity of our clusters by examining the expression of canonical ‘cell-type’ markers (i.e CD3D for T cells, CD79A for B cells), that were conserved across conditions (Figure 2D; Supplementary Figure 2).

Having aligned the datasets, we next sought to compare how PBMCs vary in response to IFN-β.As both conditions were drawn from the same pool of cells, we observed a strikingly similar proportional representation of all clusters in stimulated and control experiments (R=0.997; Figure 2E), nor did we detect any “dataset-specific” cells. However, each cell type exhibited significant changes upon IFN-β stimulation. Applying single cell differential expression tests separately for each cluster, we were able to identify constitutive markers of the IFN-β response induced in all cells (ISG15, IFIT1), as well as components of the IFN-β response that varied across cell types (i.e CXCL10 was activated primarily in myeloid cells upon stimulation) (Figure 2D). We noted that even canonical cell-type markers such as CD14 were differentially expressed by monocytes (1.98-fold down-regulation; Supplementary Figure 2) in response to stimulation, highlighting the value of our unsupervised analyses in initially classifying cells.

Focusing in on the novel subsets we were able to resolve, we compared the IFN-β response program between naïve and memory CD4+ T cells, and observed nearly identical response signatures (Supplementary Figure 3A). However, while we observed a general correlation between pDC and DC responses, we also saw stark differences that reproduced across patients (Figure 2F). When comparing the IFN-β responses across all cell types, we observed that myeloid and lymphoid cells strongly clustered together, but pDC exhibited a distinct response to IFN-β and clustered separately (Figure 2G; Supplementary Figure 3B). Therefore, in a single unsupervised analysis, our alignment procedure uncovered new cell states through integrated clustering, and allowed for the identification cell-type specific response modules that are likely to play important roles *in-vivo* during immune response to infection.

**Figure 3.**
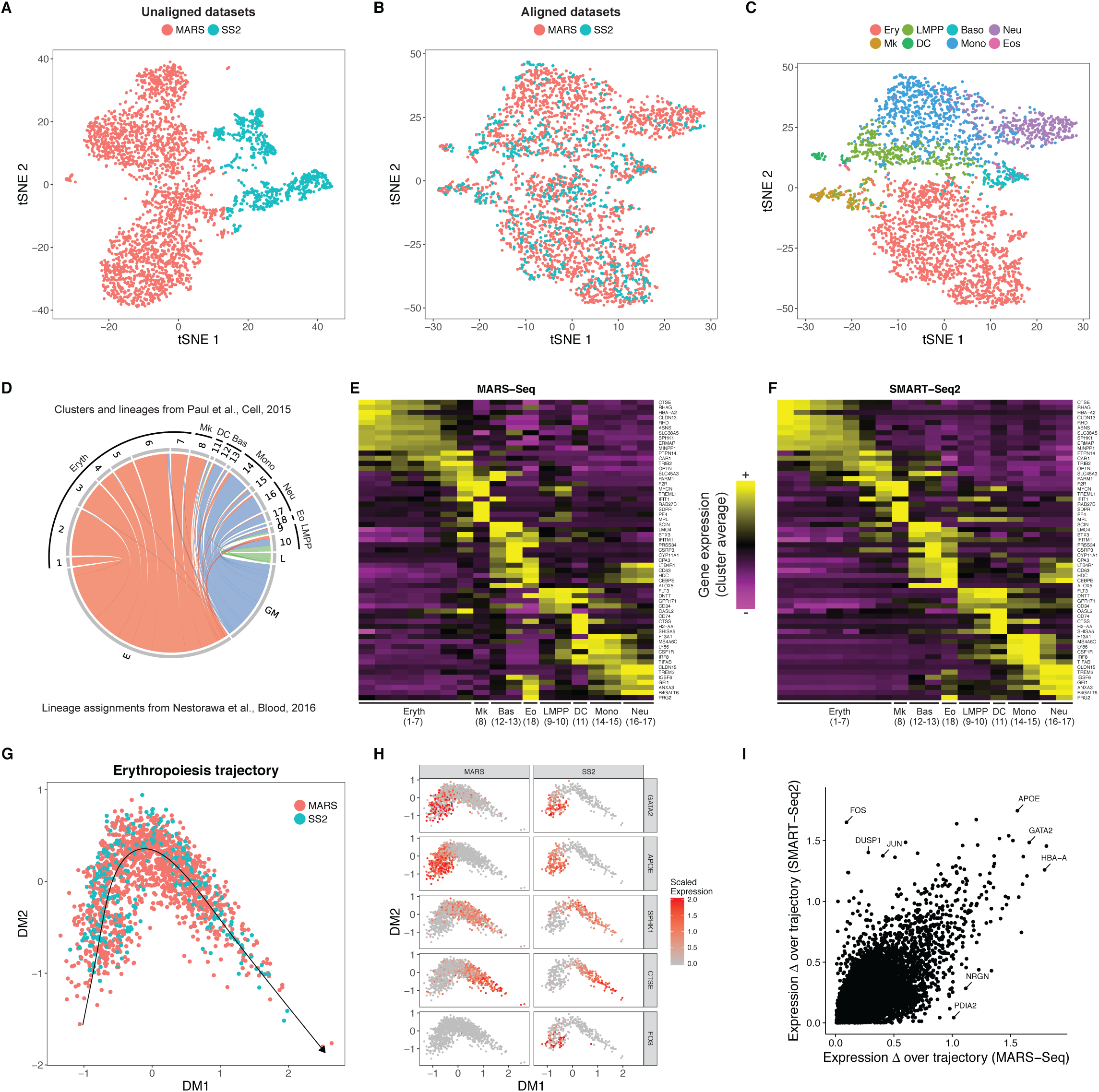
Comparative analysis of mouse hematopoietic progenitors across scRNA-seq technologies. **(A-C)** tSNE plots of 3,454 hematopoietic progenitor cells from murine bone marrow sequenced using MARS-seq (2,689) and SMART-Seq2 (765), prior to (A) and post (B-C) alignment. After alignment, cells group together based on shared progenitor type irrespective of sequencing technology. **(C-D)** Cells from the SMART-Seq2 dataset were mapped onto the closest MARS-Seq cluster and associated lineage (from Paul et al.). (C) tSNE plot of cells colored by assigned lineage. (D) Mapping correspondence between SMART-Seq2 lineage assignments (from Nestorawa et al.) and MARS-Seq clusters. **(E-F)** Heatmaps showing lineage-specific gene expression patterns in MARS-Seq and SMART-Seq2 datasets. Each column represents average expression after cells are grouped either by the original MARS-Seq cluster assignments (E), or the MARS-Seq cluster they map to (F).**(G-H)** Integrated diffusion maps of erythroid-committed cells in both datasets reveals an aligned developmental trajectory (G), with conserved ‘pseudo-temporal’ dynamics (H). **(I)** Scatter plot comparing the range in expression (absolute value) over the developmental trajectory, for each gene, across both datasets

### Integrated analysis of multiple scRNA-seq technologies

We next examined two recent single cell RNA-seq profiles of hematopoietic progenitors from murine bone marrow, but produced with starkly different technologies. Nestorowa et al.^33^ used the full-length SMART-Seq2 with deep sequencing (6,558 genes/cell) to profile 774 progenitors, while Paul et al.^34^ applied the 3’ MARS-Seq protocol with shallow sequencing (1,453 genes/cell), to examine 2,730 cells. The stark differences in amplification, normalization, and coverage pose challenges to integrate these datasets. Additionally, independent analyses from both papers highlighted different aspects of the data; the SMART-Seq2 analysis focused on the broad and continuous trajectories of cells committing to lymphoid, myeloid, erythroid lineages, while the MARS-Seq dataset identified 18 distinct clusters (and one contaminating group of NK cells), representing progenitors of eight distinct hematopoietic lineages. Despite these differences, we asked whether the same distinct progenitor subsets might be found in both datasets through integrated analysis.

Seurat alignment returned canonical correlation vectors that separated distinct progenitor subtypes, revealing populations committed to all eight distinct hematopoietic lineages in both datasets, but successfully discarded the contaminating NK population as ‘dataset-specific’ (Figure 3A-C; Supplementary Figure 4). After alignment, we mapped cells from the SMART-Seq2 dataset onto their closest cluster in the MARS-Seq dataset (Figure 3C-F; Supplementary Methods; Supplementary Table 2). We observed that early megakaryocyte-erythrocyte progenitor cells, identified in the original SMART-Seq2 publication, mapped exclusively onto erythroid and megakaryocytic progenitors in the MARS-Seq data (Clusters (C)1-7). Similarly, SMART-Seq2 granulocyte-macrophage progenitors mapped onto basophil, eosinophil, dendritic cell, neutrophil, and monocyte progenitors (C11-18). While the MARS-Seq data specifically enriched for myeloid cells, the authors identified populations of very early progenitors that were FLT3+ (C9-10). These cells represent lympho-myeloid component progenitors (lymphoid-primed multi-potent progenitors (LMPP))^35^, and early lymphoid progenitors from the SMART-Seq2 data mapped exclusively to these clusters. Indeed, after mapping, we observed nearly identical segregation of gene expression markers between SMART-Seq2 and MARS-Seq datasets (Figure 3E-F, Supplementary Figure 5), demonstrating that the biological drivers of alignment were lineage-determining factors. Therefore, Seurat alignment demonstrated that distinct committed progenitor populations were present in the SMART-Seq2 dataset, but were challenging to detect in the original analysis due to reduced cell number.

**Figure 4.**
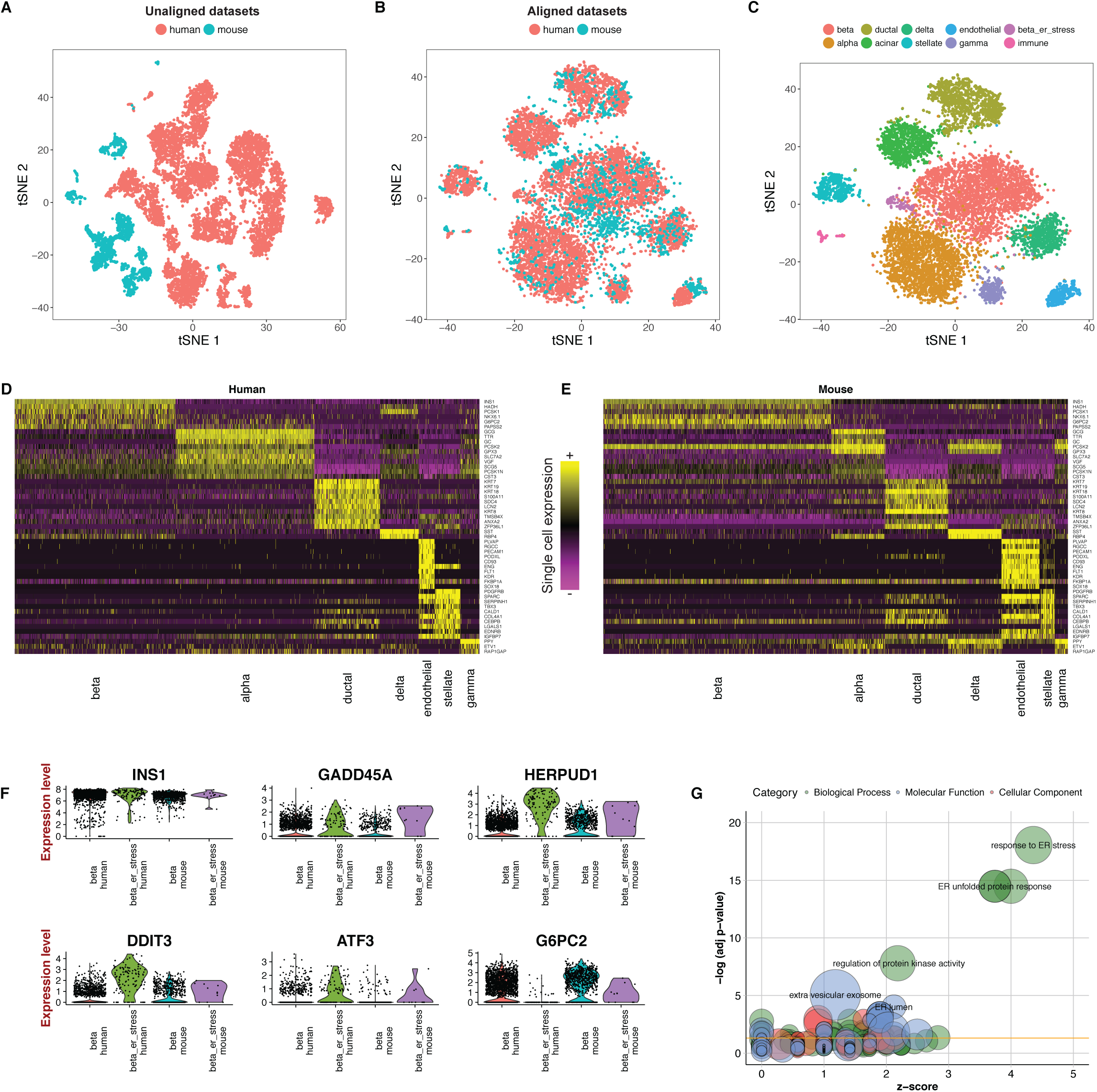
Joint identification of cell types across human and mouse islet scRNA-seq atlases. **(A-C)** tSNE plots of 10,322 pancreatic islet cells from human (8,533) and mouse (1,789) donors, prior to (A) and post (B) alignment. After alignment, cells group across species based on shared cell type, allowing for a joint clustering (C) to detect 10 cell populations. **(D-E)** Unsupervised identification of shared cell-type markers between human and mouse. Single cell expression heatmap for genes identified with joint DE testing across species. **(F)** Violin plots showing the distribution of gene expression of select genes in the beta cell and stressed beta cell clusters for both species. **(G)** Genes up-regulated in the ‘ER-stress’ subpopulation of beta cells in both species are strongly enriched for components of the ER unfolded protein stress response. GO enrichment is visualized using the GOplot R Package

**Figure 5.**
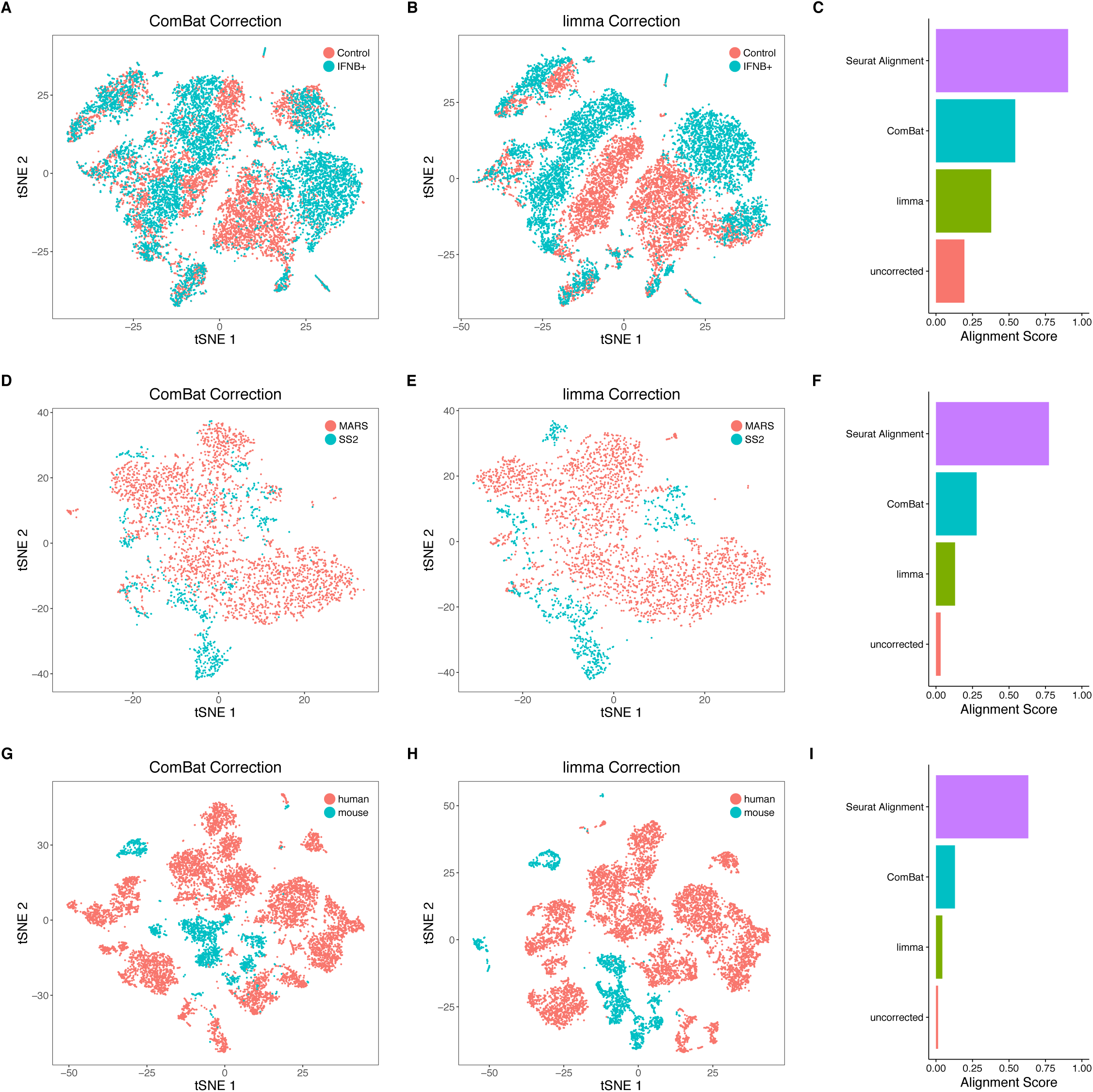
Benchmarking alignment and batch correction methods. **(A, D, G)** tSNE plots for the PBMC dataset (A), hematopoietic progenitor cell dataset (B), and pancreatic islet cell dataset (C) after correction with ComBat and **(B, E, H)** with limma. **(C, F, I)** Bar plots of the alignment score after correction using the Seurat alignment procedure, ComBat, limma, and after no correction. Seurat alignment outperforms other methods in all three demonstrated dataset pairs. Additional examples of ‘negative controls’ where Seurat fails to align datasets from different tissues are shown in Supplementary Figure 11.

Lastly, as both datasets identified developmentally heterogeneous populations during erythroid differentiation (broken into 7 stages in the MARS-Seq analysis), we applied diffusion maps to erythroid-committed cells to reconstruct a joint developmental trajectory (Figure 3G). We observed that this developmental path maintained the ‘pseudotemporal’ ordering of cells within both datasets (Supplementary Figure 6), but also aligned the two together, exhibiting nearly identical expression dynamics for canonical differentiation markers (Figure 3H). Extending this analysis globally, we observed that gene expression changes across the trajectory were largely conserved between datasets, particularly for well-characterized effectors of erythropoiesis, yet we also saw technology-specific effects -- for example a strong JUN/FOS response that has previously been associated with cellular stress during scRNA-seq^36^ (Figure 3H-I). Therefore, our procedure can successfully align both discrete and transitioning populations, and enable the identification of gene-expression programs that are conserved or unique to individual datasets.

### Joint learning of cell types across species

As a final example, we tested the ability of Seurat to align heterogeneous populations from the same tissue, but originating from different species. We examined a recent single cell study of both human and mouse pancreatic islets, performed with the inDrop technology^1^, that identified islet cell types independently in both species^37^. The study found that cell-type transcriptomes were poorly conserved between human and mouse (avg. correlation between bulk transcriptomes of individual cell types: R=0.42), often finding very few strongly expressed markers that were preserved between species. This widespread divergence poses significant challenges for integration, as structure in the dataset was largely driven by species, as well as by individual donor (Figure 4A, Supplementary Figure 7). However, we reasoned that a subset of gene-gene correlations should still be conserved, and therefore aligned all human cells against all mouse cells.

Indeed, Seurat alignment identified canonical correlation vectors that separated cell types, excluding primarily small populations of immune cells (human mast cells and murine B cells) as dataset specific (Figure 4B, Supplementary Figure 8). We next performed a single integrated clustering analysis, identifying 10 clusters, corresponding to alpha, delta, gamma, acinar, stellate, ductal, epithelial, immune, and two subgroups of beta cells (Figure 4C; Supplementary Table 3, Supplementary Figure 9). Our clusters agreed overwhelmingly with the analyses from the independent datasets^37^ (Supplementary Figure 9), though we did observe a low rate (4.3%) of discordant calls, particularly for cells with low UMI counts (Supplementary Figure 10A-B). We next designed an unbiased strategy to detect conserved cell-type specific markers across species. We first performed individual ‘within-species’ differential expression tests, and then combined the results using Fisher’s test criterion. Figure 4D-E shows the strongest shared markers of cell type using this joint differential expression analysis.

Interestingly, our fully unsupervised procedure identified a rare subpopulation of beta cells in both human and mouse. These cells expressed identical levels of INS1, but up-regulated the expression of ER stress genes (HERPUD1; GADD45A) in both species (Figure 4F). A similar signal was observed in a semi-supervised analysis of the human beta cells in the original manuscript^37^, but could not be detected in unbiased clustering, or an independent analysis of the murine dataset. In contrast, our integrated analyses reveal a conserved set of markers that are strikingly enriched for regulators of ER stress response to unfolded proteins^38,39^ (Figure 4G), which has been shown to play an important role in the onset and progression of diabetes. Notably, expression of the transcription factors ATF3 and ATF4 was highly up-regulated in both species, representing factors that have well-established roles in the initiation of stress responses in the pancreas^40,41^. Taken together, these results demonstrate that our alignment procedure can identify shared cell states even in the face of significant global transcriptional shifts, driven in this case by millions of years of evolution.

### Benchmarking alignment and batch correction techniques

We next compared Seurat’s performance to widely used batch correction tools that have been applied to both bulk^42^ and single cell genomics data^43^. To evaluate each technique, we designed an ‘alignment score’, which examines the local neighborhood of each cell after alignment (Supplementary Methods). When datasets are well aligned, this local neighborhood will consist equally of cells from both datasets, enabling us to quantify the success of each procedure with a score ranging from 0 to 1.

On the three datasets presented here, we benchmarked Seurat’s performance against ComBat^44^ and limma^45^ (Figure 5A-I). In each case, as can be visualized by tSNE or quantified with our alignment score, Seurat’s integration procedure yielded superior results. The differences between these procedures was particularly striking when the transcriptomic differences between datasets (i.e ‘batch effect’) significantly outweighed differences between cell types (‘biology’), as in cross-species integration. However, when we attempted to align datasets from different tissues as a negative control, we observed poor results and low alignment scores, even when cells are not automatically classified as ‘dataset-specific’ (Supplementary Figure 11).

## DISCUSSION

We developed a strategy to integrate scRNA-seq datasets by identifying shared sources of variation between two datasets, corresponding to subpopulations present in both experiments. Implemented in our R toolkit Seurat, our procedure tackles several technical challenges, including the unbiased identification of shared gene-gene correlations across datasets, as well as the alignment of canonical correlation vectors using non-linear ‘warping’ algorithms. Below, we briefly discuss the potential utility and future challenges for these and similar methods.

Dataset integration represents a key step in a general framework for case/control studies performed with single cell resolution. Our analysis of IFN-β stimulated and control PBMCs enabled us to discover how each of 13 separate hematopoietic populations responded to IFN-β stimulation. The ability to reveal cell-type specific responses through unsupervised analysis exemplifies the potential of scRNA-seq to provide new levels of resolution to comparative analyses. As new datasets are generated, we expect that similar computational analyses will be invaluable for characterizing the immune system’s response to vaccination, inflammatory disease, and cancer. Extending beyond environmental perturbations, integrated analysis will also provide deeper insight into how genetic variation and manipulation affect heterogeneous populations.

Recent benchmarking studies of diverse scRNA-seq^46,47^ technologies have consistently demonstrated that no single method is uniformly superior, but rather, that each has individual strengths and weaknesses. Here, we demonstrate that integrating scRNA-seq data produced with technologies that emphasize both ‘breadth’ (emphasizing cell number over sequencing depth), and ‘depth’ (prioritizing deep sequencing per cell), can combine the benefits of both approaches. As further demonstration, we provide an additional alignment example in the form of an R Markdown tutorial (Supplementary Note 1), demonstrating how to align human PBMC datasets produced with the 10X Chromium System, and SeqWell^48^. We anticipate that these methods will enable new analyses for a diverse set of labs, as well as for consortia such as the Human Cell Atlas^15,37,49^, which aims to define all human cell types by integrating data generated across diverse single cell ‘omics’ approaches.

Lastly, we demonstrate the ability to align differentiated cell types between human and mouse pancreatic islets, identifying a shared population of beta cells responding to ER protein misfolding stress. These and similar analyses may provide invaluable comparative tools for studies utilizing mouse models of human disease, potentially enabling the identification of human correlates of pathogenic populations discovered in mouse (or vice versa). Furthermore, new datasets will enable the alignment and comparison of developmental trajectories across species, leading to a deeper understanding of how the gene regulatory networks generating cellular diversity are rewired across evolution.

While Seurat’s current approach performs pairwise alignment of two datasets, these approaches can be generalized to multiple datasets either by repeated pairwise alignment to a reference, or through extensions of CCA to analyze multiple samples (i.e. multi-set CCA)^50,51^. Additionally, while we focus here on alignment of sequencing-based datasets, the recent invention of spatially-resolved or *in-situ* methods for transcriptomic profiling^52-54^ raise the exciting potential for integration with scRNA-seq datasets, including the ability to extend previous efforts to spatially resolve scRNA-seq data^9,10^ towards an unsupervised procedure generalizable to any tissue. While in a nascent stage compared to bulk methods, we therefore anticipate the continued development of exciting and impactful strategies for integrated analysis of single cell ‘omics’.

## ACKNOWLEDGEMENTS

We thank members of the Satija lab, as well as G. Fishell, C. Desplan, R. Bonneau, and E. Macosko for valuable feedback, and F. Hamey, HM Kang, and J. Ye for assistance with published datasets. This work was supported by an NIH New Innovator Award (1DP2HG009623-01) to RS and an NSF Graduate Fellowship (DGE1342536) to AB. Any opinions, findings, and conclusions or recommendations expressed in this material are those of the author(s) and do not necessarily reflect the views of the National Science Foundation.

## AUTHOR CONTRIBUTIONS

AB and RS conceived the research, implemented the alignment procedure, performed all data analysis, and wrote the manuscript.

